# Intermolecular Energy Migration via HomoFRET Captures the Modulation in the Material Property of Phase-Separated Biomolecular Condensates

**DOI:** 10.1101/2024.02.12.579899

**Authors:** Ashish Joshi, Anuja Walimbe, Snehasis Sarkar, Lisha Arora, Gaganpreet Kaur, Prince Jhandai, Dhruba Chatterjee, Indranil Banerjee, Samrat Mukhopadhyay

## Abstract

Biomolecular condensation via phase separation of proteins and nucleic acids has emerged as a crucial mechanism underlying the spatiotemporal organization of cellular components into functional membraneless organelles. However, aberrant maturation of these dynamic, liquid-like assemblies into irreversible gel-like or solid-like aggregates is associated with a wide range of fatal neurodegenerative diseases. New tools are essential to dissect the changes in the internal material properties of these biomolecular condensates that are often modulated by a wide range of factors involving the sequence composition, truncations, mutations, post-translational modifications, and the stoichiometry of nucleic acids and other biomolecules. Here, we employ homo-Förster Resonance Energy Transfer (homoFRET) as a proximity ruler to study intermolecular energy migration that illuminates the molecular packing in the nanometric length-scale within biomolecular condensates. We used the homoFRET efficiency, measured by a loss in the fluorescence anisotropy due to rapid depolarization, as a readout of the molecular packing giving rise to material properties of biomolecular condensates. Using single-droplet anisotropy imaging, we recorded spatially-resolved homoFRET efficiencies of condensates formed by fluorescent protein-tagged Fused in Sarcoma (FUS). By performing single-droplet picosecond time-resolved anisotropy measurements, we were able to discern various energy migration events within the dense network of polypeptide chains in FUS condensates. Our homoFRET studies also captured the modulation of material properties by RNA, ATP, and post-translational modification. Additionally, we utilized mammalian cell lines stably expressing FUS to study nuclear FUS and oxidative stress-induced stress granule formation in the cytoplasm. Our studies demonstrate that spatially-resolved homoFRET methodology offers a potent tool for studying intracellular phase transitions in cell physiology and disease.

## Introduction

Biomolecular condensation involves the formation of dynamic, liquid-like assemblies via the phase separation of proteins in association with nucleic acids. These membraneless organelles or MLOs achieve cellular compartmentalization and facilitate the spatiotemporal organization of an array of critical cellular processes across organisms^1–10^. The class of proteins termed intrinsically disordered proteins or regions (IDPs/IDRs) with or without nucleic acids form dynamic, liquid-like, selectively permeable, non-stoichiometric assemblies such as stress granules, germ granules, P bodies, Cajal bodies, nuclear paraspeckles, etc., which are closely associated with critical cellular functions, including cellular signaling, regulation, genome organization, transcription, translation, and immune responses^6,8,11–18^. These tightly regulated assemblies, when dysregulated, lead to the formation of irreversible solid-like aggregates associated with several neuropathological disorders, including Alzheimer’s disease, amyotrophic lateral sclerosis (ALS), frontotemporal lobar degeneration (FTLD), and so on^14,15,19,20^. Thus, the need to uncover the molecular drivers that govern the fate of these supramolecular assemblies by modulating their material properties has led to the development of a multitude of techniques for investigating their diffusion characteristics, viscoelasticity, molecular packaging, nanoscale organization, and so on. Several microscopic techniques like fluorescence recovery after photobleaching (FRAP), fluorescence loss in photobleaching (FLIP), atomic force microscopy (AFM), etc., have been adapted to probe condensate properties, such as morphology, fusion, size distribution, and diffusion properties^21,22^. Additionally, certain microfluidic-based approaches, along with passive microrheology, monitor the diffusion of beads within the condensates or dense phase. These techniques can be successfully performed within condensates of significantly larger sizes or the ensemble condensed phase after the careful selection and processing of the beads^21,22^. However, majority of the aforementioned methodologies lack non-invasive, broader application and the ability to illuminate the condensate organization at the nanoscale, crucial in dictating the mesoscopic properties and the fate of condensates.

Similarly, Förster Resonance Energy Transfer (FRET) is an alternative technique to study the protein-protein interactions quantitively^23–25^. The typically used heteroFRET utilizes two FRET-compatible fluorophores covalently linked at specific positions to the protein chain, which proves a cumbersome task, especially when performing cellular studies. On the contrary, tagging the proteins of interest with fluorescent proteins is an extensively used methodology for studies investigating the expression, localization, functioning, and protein-protein interactions within cells. Along these lines, homoFRET, a special case of FRET, utilizes a single fluorophore with a small Stokes’ shift and a significant overlap between its fluorescence excitation and emission spectrum and has been previously captured in the form of fluorescence depolarization or loss in anisotropy as a function of local fluorophore densities^26–30^. Thus, homoFRET can be employed to investigate the proximal and distal molecular packaging within mesoscopic supramolecular assemblies such as phase-separated biomolecular condensates. Here, we demonstrate the application of a simple and versatile technique of anisotropy imaging to study homoFRET as a readout for the internal architecture and supramolecular packing governed by the intermolecular interactions and nanoscale clustering within the dynamic biomolecular condensates. HomoFRET imaging captures the enhanced excited state energy migration indicating the reduced intermolecular distances and densely-packed organization within the condensates of an archetypal phase-separating protein Fused in Sarcoma (FUS). In conjunction with anisotropy imaging, our picosecond time-resolved anisotropy measurements allow us to discern the components of fluorescence depolarization originating from diverse modes of energy migration. Our steady-state anisotropy imaging also sheds light on the role of small-molecule phase separation modulators such as RNA and ATP, along with post-translational modifications in altering the protein-protein associations and thus, the molecular packaging within these dynamic assemblies of FUS. The anisotropy decay kinetics and corresponding time constants obtained within these droplets further illuminate molecular events and condensate properties responsible for facilitating energy migration via homoFRET. Lastly, we also show the utility of anisotropy imaging within phase-separated assemblies of FUS formed *in situ*, under diverse cellular conditions. The anisotropy imaging tool to detect homoFRET in condensates can provide a highly sensitive and potent tool for the preliminary detection of the diverse complex biomolecular condensates formed *in vitro* and within cells and further illuminate their internal molecular organization and packaging.

## Results

### Mechanism and detection of excitation energy migration via HomoFRET

HomoFRET can be defined as energy migration from the excited state of a donor fluorophore to an acceptor fluorophore of similar chemical identity when two or more of them are placed proximally (Fig. 1a). Excitation energy migration via homoFRET can be observed in the case of fluorophores with a small Stokes’ shift and a significant overlap between the excitation and emission spectrum. HomoFRET involves energy transfer in a non-radiative but reversible manner, giving rise to a donor lifetime indistinguishable from the donor lifetime obtained in the absence of energy transfer. Hence, unlike heteroFRET, homoFRET cannot be detected by measuring a change in the donor lifetime as we observe an identical acceptor molecule in the same spectral region. Excitation with polarized light leads to similarly polarized emission in the case of zero energy transfer. However, in the case of homoFRET, energy migration to a fluorophore with a non-identically oriented dipole leads to a rapid depolarization of the emitted fluorescence. This fluorescence depolarization or loss in anisotropy can be easily captured by anisotropy imaging, where the overall extent of depolarization is determined by the local crowding or packing and is independent of the rotational diffusion of the fluorophore molecules (Fig. 1a). This loss in anisotropy due to homoFRET is accompanied by the introduction of an additional faster component in the fluorescence anisotropy decay. This faster component is absent in the anisotropy decay observed under non-homoFRET conditions, provided by time-resolved fluorescence anisotropy measurements. The class of fluorescent proteins comprising green fluorescent protein (GFP) and its genetically encoded variants, including the yellow fluorescent protein (YFP), blue fluorescent protein (BFP), cyan fluorescent protein (CFP), etc., can serve as suitable candidates for homoFRET, owing to their much slower rotational mobility and thus high intrinsic anisotropy due to their bulky volume. Here, we used the enhanced green fluorescent protein (eGFP) as a homoFRET reporter and studied FUS tagged with eGFP in both the mixed (dispersed) and demixed (condensed) phases using fluorescence anisotropy imaging and picosecond time-resolved anisotropy measurements. Our microscopy setup consists of a polarized excitation source (485 nm pulsed laser), an inverted microscope, and an integrated detection system comprising a dichroic mirror, bandpass filter, pinhole, polarizing beamsplitter, and detectors with single-photon detection (single-photon avalanche diodes; SPADs) (Fig. 1b). For anisotropy imaging, the emitted fluorescence is separated into parallel and perpendicular channels, which are then used to construct images after incorporating the respective correction factors for the objective lens and differential detector efficiencies. Utilizing this setup, we began with our anisotropy imaging to explore the molecular packaging and organization within individual condensates of FUS formed under different phase separation conditions, in a droplet-by-droplet manner. The pulsed laser, in combination with time-gated single-photon detection, allowed us to perform picosecond time-resolved fluorescence anisotropy measurements within single droplets to resolve and capture the distinct energy migration rates or time constants, which otherwise remain inaccessible in the time-averaged steady-state measurements. Lastly, using this setup, we extended this technique of anisotropy imaging to detect and investigate the nuclear and cytoplasmic liquid-like assemblies or puncta of FUS-eGFP formed *in situ* in a puncta-by-puncta manner.

**Fig. 1.**
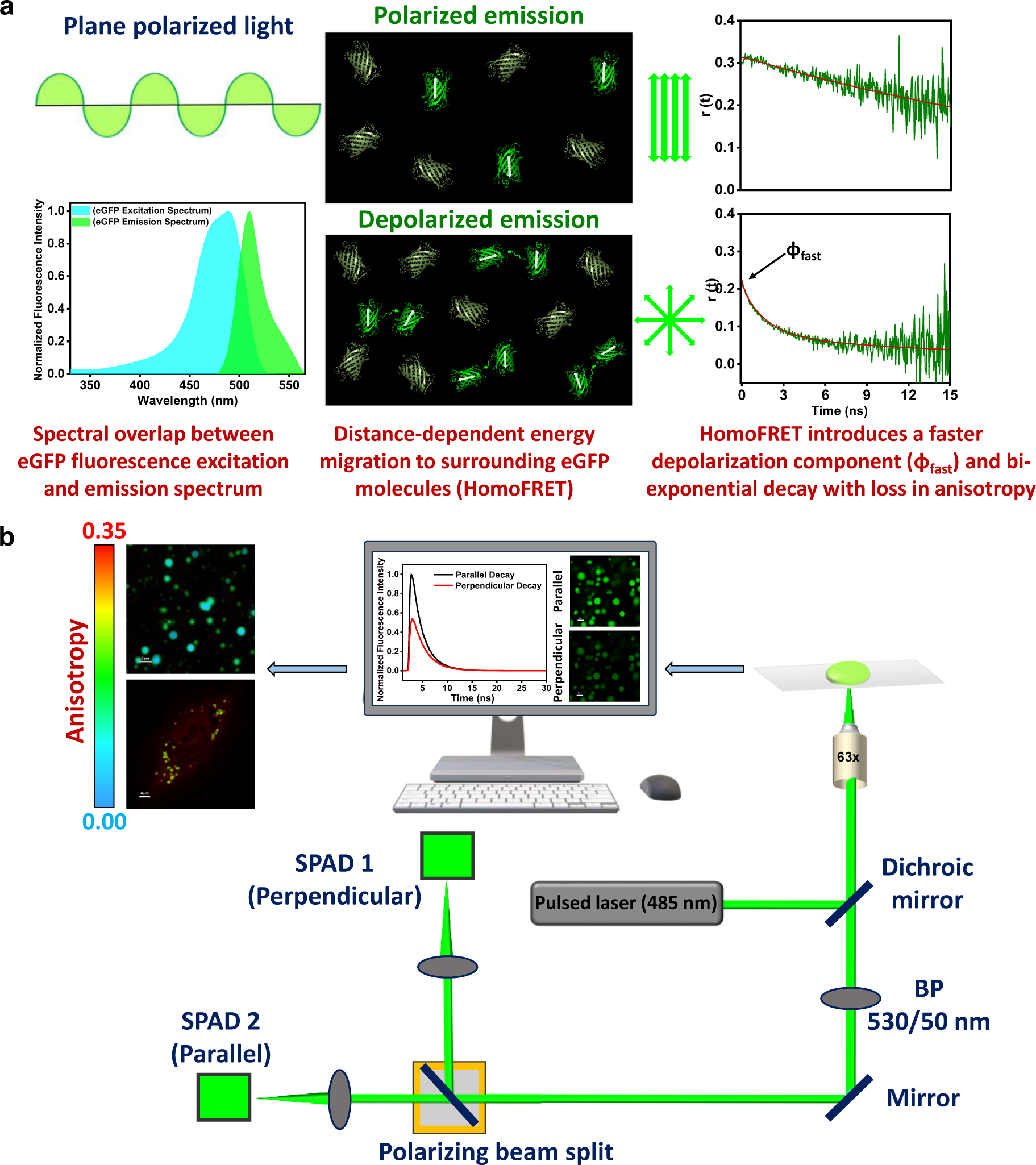
Schematic illustration depicting the mechanism and detection of homoFRET. a. Principle of energy migration via homoFRET. Fluorescent proteins with spectral overlap between their excitation and emission spectrum can exhibit a non-radiative excitation energy migration via homoFRET. At larger fluorophore concentrations and smaller intermolecular distances, the transfer of energy to differently oriented fluorophores in proximity leads to a depolarization of the emitted fluorescence. The magnitude of anisotropy loss indicates the extent of homoFRET and, indirectly, the local fluorophore densities. HomoFRET introduces a faster decay component in time domain anisotropy measurements. b. Confocal microscopy setup (MicroTime 200, PicoQuant) used for our single-droplet anisotropy imaging and picosecond time-resolved measurements. Our setup constitutes a pulsed laser (485 nm), an inverted microscope, and a main optical unit comprising a dichroic mirror, pinhole, polarizing beamsplitter, and avalanche photodiodes with single-photon detection efficiency. After filtering out the out-of-focus light by pinhole, the emitted fluorescence is further separated and directed into the parallel and perpendicular channels (SPADs) by the polarizing beamsplitter. See “Methods” for more details.

### HomoFRET imaging as a proximity ruler for biomolecular condensation

FUS comprises a multidomain architecture with a low-complexity (LC) prion-like N-terminal domain and a partially structured RNA-binding C-terminal domain (Fig. 2a)^31,32^. In order to probe phase separation of FUS via homoFRET, we utilized C-terminally eGFP tagged FUS (MBP-tev-FUS-eGFP) for our homoFRET measurements. As a prelude, we began with anisotropy imaging in the dispersed phase and condensed phases of FUS to obtain steady-state anisotropy values. In the monomeric condition (200 nM FUS-eGFP), we obtained a higher anisotropy value corresponding to the absence of homoFRET due to the low fluorophore densities in the lightly packed dispersed state (Fig. 4.2b, c). Next, to monitor the condensed phase, droplet formation was initiated by the addition of TEV protease for cleavage of the solubilizing MBP tag. Upon phase separation, we obtained a large dip in the steady-state anisotropy value within individual droplets, indicating higher excited state energy migration due to the dense, closely-packed organization of FUS-eGFP molecules, as expected, within the condensed phase (Fig. 2b, c). This increase in the local density of the FUS-eGFP molecules could be quantified by estimating the energy migration or homoFRET efficiency (E_homoFRET_) based on the relative change in steady-state anisotropy values from the dispersed (non-homoFRET) to condensed (homoFRET) phases, or the loss in steady-state fluorescence anisotropy value due to homoFRET (Δr_0_). We could calculate an apparent energy migration efficiency (E_homoFRET_) of ∼ 0.66 within the condensates of FUS-eGFP (Fig. 2d); see “Methods” for more details. Next, to validate that this drop in fluorescence anisotropy was indeed a consequence of the increased homoFRET within condensates, we subjected these droplets to photobleaching with a high-intensity laser for a longer duration and monitored the fluorescence intensity for recovery of anisotropy value with time. As expected, we obtained a gradual recovery of the anisotropy values within the condensates to that of non-homoFRET conditions upon bleaching the samples (Fig. 2e). Next, to probe the depolarization kinetics originating from homoFRET, we set out to perform picosecond time-resolved anisotropy measurements. In the monomeric phase, the anisotropy decay followed a typical monoexponential kinetics with a fundamental anisotropy (r_0_ ∼ 0.31) and a correlation time (ϕ) of ∼ 30 ns, corresponding to the rotational dynamics of FUS-eGFP in the dispersed state (Fig. 2f). On the contrary, we obtained three different components of anisotropy depolarization from the decay kinetics within the droplet phase. The ultrafast component presented in the form of a loss in the fundamental anisotropy (τ_EM1_ with amplitude β_EM1_ ∼ 0.26), where the rate of energy migration was much faster (subnanosecond) than our experimental temporal resolution. The anisotropy decay profile exhibited characteristic biexponential decay kinetics with an intermediate time constant (τ_EM2_) of ∼ 1.5 ns and a slow time constant (τ_EM3_) of ∼ 25 ns (Fig. 2f). These nanosecond time constants represent the intermediate and slower rates of excitation energy migration, which in association with the ultrafast subnanosecond component, collectively contribute toward the emission depolarization captured in our homoFRET measurements.

**Fig. 2.**
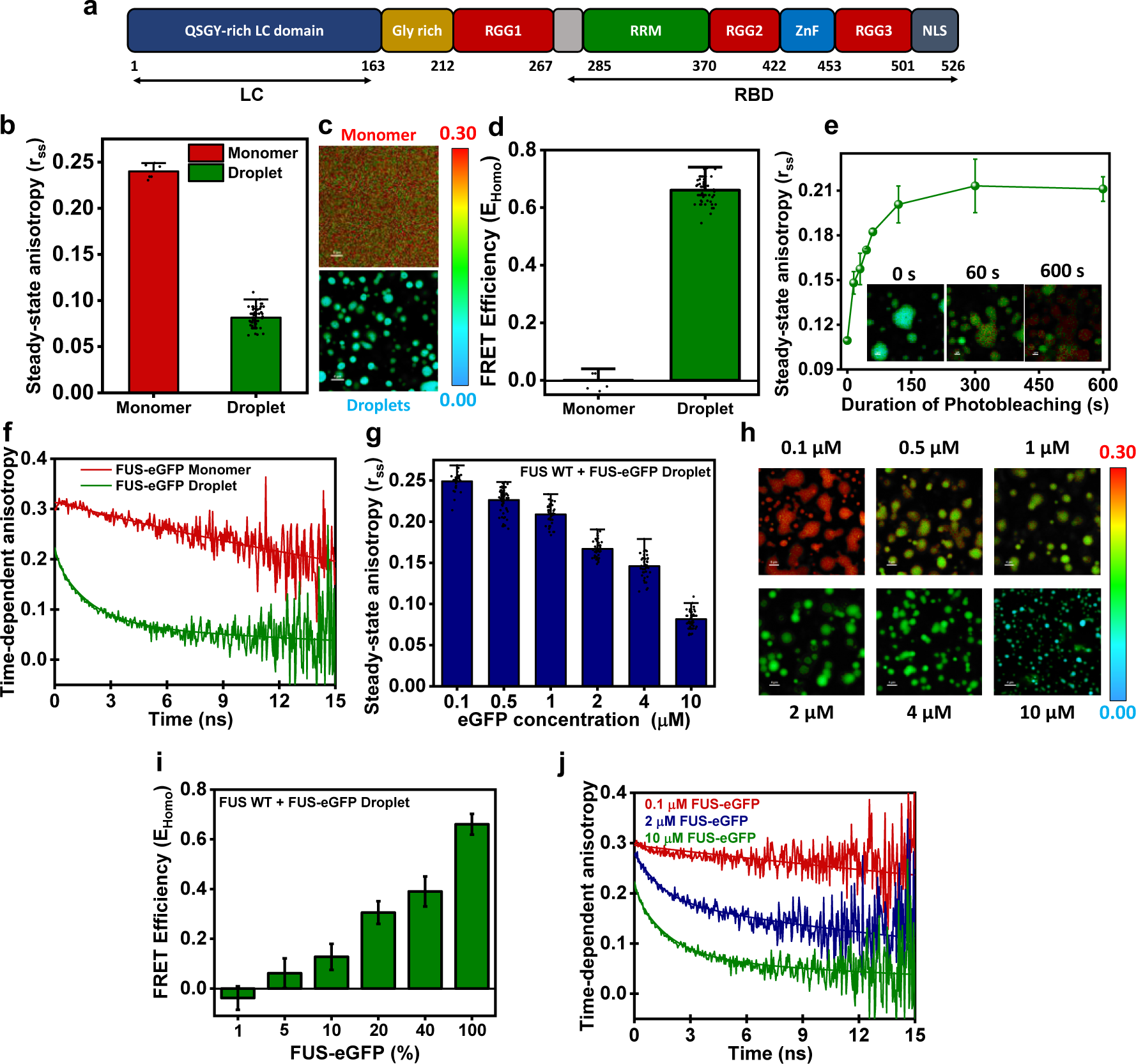
HomoFRET within phase-separated FUS-eGFP condensates. a. Sequence architecture of Fused in Sarcoma (FUS) depicting the N-terminal SYGQ-rich low-complexity domain (LC), the Arg-Gly rich RGG domains, an RNA-recognition motif (RRM), a zinc-finger domain (ZnF) and a C-terminal nuclear localization signal (NLS). b. Steady-state anisotropy values and c. the representative anisotropy images obtained from the monomeric phase and single droplets of FUS-eGFP. d. The calculated FRET efficiencies obtained from steady-state anisotropy values indicate the extent of molecular packaging in the dispersed phase and single droplets. Data represent mean ± SD (n = 6 and 60 for monomeric samples and droplets, respectively). e. Steady-state anisotropy values and their representative images (inset) obtained at high laser intensity as a function of time. Data represent mean ± SD (n = 3 independent reactions). f. Picosecond time-resolved fluorescence anisotropy decay obtained in the monomeric phase, and single droplets of FUS-eGFP fitted using monoexponential decay kinetics (no homoFRET) and a biexponential decay kinetics (high-homoFRET), respectively. g. Steady-state anisotropy values, their representative anisotropy images (h), and corresponding homoFRET efficiencies (i) as a function of varying FUS-eGFP concentration within FUS droplets. Data represent mean ± SD (n ≥ 30 for single droplets). j. Representative time-resolved anisotropy decay at varying fluorophore concentrations yielded monoexponential decay kinetics (100 nM FUS-eGFP) and biexponential decay kinetics (2 µM and 10 µM FUS-eGFP) with individual energy migration time constants. Rotational correlation time and energy migration time constants are included in Table 1.

**Table 1.** Excitation energy migration time constants and rotational correlation times with associated amplitudes recovered by fitting fluorescence anisotropy decay kinetics obtained from time-resolved measurements.

|  | **Ultrafast**<br>**energy**<br>**migration**<br>**time constant**<br>**(Loss in**<br>**fundamental**<br>**anisotropy)**<br>**($\beta_{EM1}$)** | **Intermediate**<br>**energy**<br>**migration**<br>**time constant**<br>**($\tau_{EM2}$) and**<br>**amplitude**<br>**($\beta_{EM2}$)** | **Slow energy**<br>**migration time**<br>**constant ($\tau_{EM3}$)**<br>**and amplitude**<br>**($\beta_{EM3}$)** | **Rotational**<br>**correlation time**<br>**($\phi$) and**<br>**amplitude ($\beta_\phi$)** |
| --- | --- | --- | --- | --- |
| Monomer | - | - | - | $31.34 \pm 0.82$ ns |
| 1% FUS-eGFP in<br>10 μM FUS<br>Droplet | - | - | - | $67.92 \pm 5.01$ ns |
| 20% FUS-eGFP in<br>10 μM FUS<br>Droplet | 0.06 | $1.61 \pm 0.21$ ns<br>($0.37 \pm 0.03$) | $19.86 \pm 3.11$ ns<br>($0.28 \pm 0.08$) | $58.34 \pm 12.35$ ns<br>($0.37 \pm 0.06$) |
| 10 μM FUS-eGFP<br>Droplet | 0.26 | $1.73 \pm 0.14$ ns<br>($0.65 \pm 0.04$) | $18.74 \pm 4.75$ ns<br>($0.35 \pm 0.04$) | - |
| Methylated 10 μM<br>FUS-eGFP Droplet | 0.23 | $1.11 \pm 0.11$ ns<br>($0.39 \pm 0.05$) | $9.03 \pm 1.10$ ns<br>($0.61 \pm 0.04$) | - |
| 10 mM ATP 10<br>μM FUS-eGFP<br>Droplet | 0.03 | $1.60 \pm 0.24$ ns<br>($0.16 \pm 0.03$) | - | $49.63 \pm 7.02$ ns<br>($0.84 \pm 0.03$) |

To further explore the sensitivity of homoFRET imaging towards differential intermolecular distances, we set out to perform anisotropy measurements over a wide range of fluorophore densities by varying the fraction of FUS-eGFP within the condensates of FUS. Beginning with one percent, we increased the fraction of FUS-eGFP and captured an incremental loss in the steady-state anisotropy values (Fig. 2g, h) corresponding to the enhanced energy migration efficiencies (apparent homoFRET efficiencies, E_homoFRET_) between the progressively closely-packed FUS-eGFP molecules (Fig. 2i). At a low fluorophore concentration (1%), local densities of FUS-eGFP molecules resulted in steady-state anisotropy values reminiscent of non-homoFRET conditions, due to the large average intermolecular distances within condensates, leading to negligible or no energy migration. This absence of excitation energy migration was evident from our picosecond time-resolved measurements, which exhibited a typical monoexponential decay kinetics (Fig. 2j). The correlation time constant obtained from monoexponential fitting (∼ 65 ns) corresponds to the rotational dynamics of FUS-eGFP (ϕ) within the viscoelastic interior of FUS droplets. This slower depolarization component could be captured within these droplets as a result of the undepolarized axis left due to the absence of faster decaying homoFRET components. This slower rotational correlation time could also be captured by fitting the anisotropy decay of droplets with a 20% labeled fraction with triexponential decay kinetics. At this fraction of FUS-eGFP, we could obtain an extremely minor contribution of the ultrafast component represented by the loss in r_0_ (τ_EM1_ with amplitude β_EM1_ ∼ 0.06), an intermediate time constant (τ_EM2_, ∼ 1.6 ns), the slow time constant (τ_EM3_, ∼ 20 ns), and the slower correlation time (ϕ) corresponding to the slower rotational diffusion (∼ 60 ns) within droplets. These time constants shed light on the multiple modes of excitation energy migration originating from the array of dynamic molecular events, comprising the transient making and breaking of contacts, conformational fluctuations, translational diffusion, etc., operating at their characteristic timescales, in turn determining the rate of energy migration within condensates. These multivalent fuzzy interactions contribute to determining the mesoscopic attributes, including material properties, viscoelasticity, supramolecular packaging, and nanoscale organization within biomolecular condensates. We next asked whether homoFRET imaging can capture the effect of RNA on the droplet interior of FUS-RNA heterotypic condensates.

### Illuminating internal architecture of FUS-RNA heterotypic condensates using homoFRET imaging

In association with proteins, nucleic acids form the primary constituents of the vast majority of cellular biomolecular condensates^6,33–35^. FUS is a well-known RNA-binding protein involved in some of the critical DNA/RNA-associated cellular functions, where RNA is known to modulate the phase behavior and properties of FUS condensates^36–38^. RNA is capable of tuning the assembly, droplet interior (material properties), and dissolution of condensates of RNA-binding proteins in a concentration-dependent manner. Hence, we next set out to investigate the effect of RNA on the condensates of FUS using homoFRET imaging. Due to the high concentration and densely packed droplet interior, even in the absence of RNA, FUS-eGFP droplets exhibit extremely low anisotropy values, as shown in our previous results. Upon the addition of RNA, we captured a further dip in the anisotropy values (Fig. 3a), indicating the enhanced molecular interactions and closer packaging, in agreement with the phase separation-promoting behavior of RNA. However, due to the initially borderline values, we shifted to an intermediate fluorophore fraction to monitor the effect of a wider range of RNA concentrations and to assess the changes in fluorophore densities with higher precision. Thus, we proceeded with our RNA-dependent homoFRET measurements at a 20% fluorophore concentration doped within the heterotypic FUS:RNA condensates formed in the presence of varying concentrations of polyU RNA. With increasing RNA concentration, the anisotropy imaging showed an increase in the emission depolarization (Fig. 3b, c), originating from the enhanced excitation energy migration (Fig. 3d), presumably due to the increasingly compact packaging within FUS:RNA condensates. While facilitating and participating in multivalent protein-RNA interactions, RNA possibly enables the formation of dense intermolecular protein-protein and protein-RNA networks, leading to a reduction in the average intermolecular distances between the FUS-eGFP molecules inside condensates. This condensing effect of RNA has been well-established for multiple phase-separating RNA-binding proteins^39,40^, corroborating our FRAP measurements, which showed a slower diffusion within the densely crowded droplet interior in the presence of lower concentrations of RNA (Fig. 3e).

**Fig. 3.**
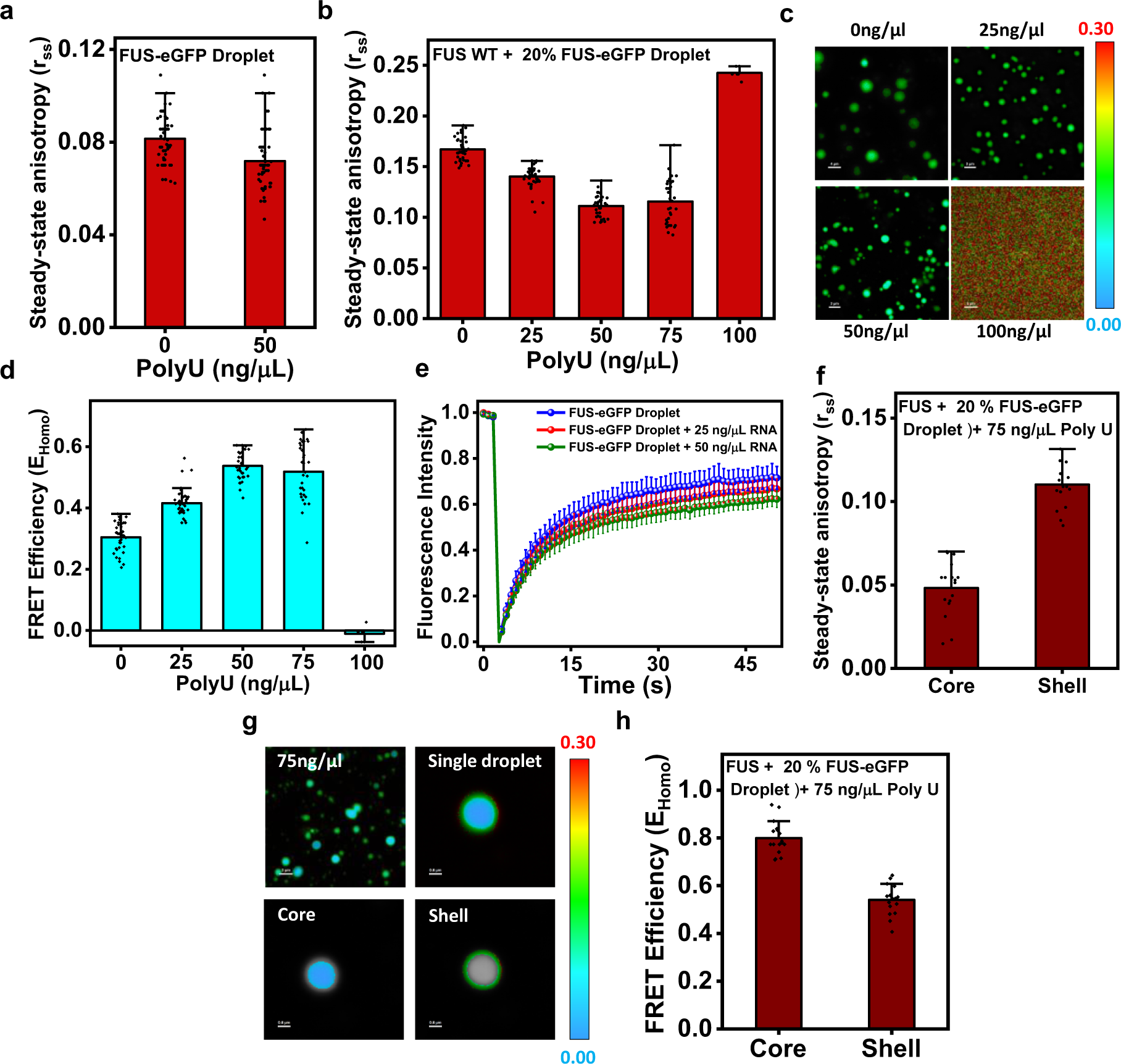
HomoFRET imaging detects nanoscale organization within FUS:RNA heterotypic condensates. a. Steady-state anisotropy plot indicating the slight dip in anisotropy within single droplets of FUS-eGFP in the presence of RNA. Data represent mean ± SD (n = 60 for single droplets). Data for 0 ng/µL RNA is shown for comparison, same as in 2b, g) b. Single-droplet steady-state anisotropy plot of FUS droplets doped with 20 % FUS-eGFP with varying RNA. Data represent mean ± SD (n ≥ 30 for 0, 25, 50, 75 ng/µL RNA and n = 6 for 100 ng/µL RNA). c. Representative anisotropy images of FUS droplets in the range of RNA concentrations plotted in (b). d. Apparent homoFRET efficiencies as a function of RNA concentrations. estimated from (b). e. Fluorescence recovery after photobleaching (FRAP) kinetics of FUS-eGFP droplets in the absence and presence of RNA. Data represent mean ± SD (n = 5, 5, and 7 for 0 ng/µL, 25 ng/µL, and 50 ng/µL RNA, respectively). f. Steady-state anisotropy plots and g. representative images showing the core-shell morphology of droplets formed at a polyU concentration of 75 ng/µL. Data represent mean ± SD (n = 17 single droplets). h. The calculated FRET efficiency values suggest a densely packed protein core enclosed within a lightly packed shell.

Upon further increasing the concentration, homoFRET imaging revealed a slight reduction followed by a complete abrogation of excitation energy migration as evident from the corresponding anisotropy values obtained for 75 and 100 ng/µL RNA, respectively (Fig. 3b, c). The lower FRET efficiency indicated a decrease in the droplet compaction, ensuing a complete dissolution of the droplets at 100 ng/µL RNA (Fig. 3d). This was reflected in the complete recovery of steady-state anisotropy to that of the non-homoFRET conditions. Interestingly, the anisotropy imaging of condensates in the presence of 75 ng/µL RNA exhibited density variations and a specialized nanoscale organization inside the majority of the droplets. The droplet core showed consistently lower anisotropy, as compared to the periphery, as visible in the representative anisotropy images (Fig. 3f, g). Upon closer examination and analysis, we obtained significantly distinct protein packaging and molecular organization as suggested by the higher energy migration efficiency at the center, as opposed to the peripheral regions of these droplets (Fig. 3h). Our homoFRET studies hint towards the existence of a typical core-shell architecture, with a comparatively denser core (high local protein densities) surrounded by a relatively lightly-packed shell inside the heterotypic FUS:RNA condensates. These peculiar droplet morphologies originate from the variable density distributions, where the densely packed core harbors a large number of protein and RNA molecules enmeshed in an extensive network of heterotypic protein-RNA interactions. These multiphasic architectures have previously been reported *in vitro* for multicomponent co-phase separation systems^41–43^. Such immiscible multiphasic morphologies originating from heterogeneous nanoscale architecture within cellular condensates can be readily captured by this robust methodology of homoFRET imaging. Taken together, our homoFRET imaging provides a readout for the local molecular clustering and the average intermolecular distances of protein molecules within the heterotypic FUS:RNA condensates. Further, we sought to probe the alterations in droplet organization within FUS-eGFP condensates as a result of other phase separation modulators, such as post-translational modification and ATP.

### HomoFRET captures the altered molecular packaging due to post-translational methylation and ATP

The partially structured C-terminal domain of FUS is enriched in amino acids arginine and glycine (RGG-rich domains) (Fig. 2a) and is identified to play an essential role in the RNA-binding activity and biomolecular condensation of FUS^33,36,37,44^. In healthy neurons, the arginine residues in the CTD of FUS are post-translationally modified by extensive methylation and can harbor either one (monomethylated Arg) or two (dimethylated Arg) methyl groups^45,46^. Several reports show that post-translational methylation is disrupted in the pathology of FTLD-FUS, leading to the formation and deposition of insoluble cytoplasmic aggregates of hypomethylated FUS^45,46^. Furthermore, methylation has been shown to alter the phase behavior and retain the liquid-like dynamics within FUS condensates, highlighting the significance of investigating the role of methylation in determining the material properties and internal organization of FUS condensates. With this objective, we next aimed to capture this change in droplet interior within FUS-eGFP condensates upon methylation. In this direction, we performed *in vitro* methylation using a previously established protocol employing S-adenosyl-methionine (SAM) as the methyl-group donor and the enzyme protein arginine methyltransferase 1 (PRMT1) to catalyze the reaction^45^, see “Methods” for more details. We began with anisotropy imaging of methylated FUS-eGFP droplets, which revealed an increased steady-state anisotropy value in comparison to the unmethylated FUS-eGFP droplets, as evident in the representative anisotropy images (Fig. 4a, b). The calculated homoFRET efficiency (Fig. 4c) showed a significant dip in the energy migration, indicating a reduced protein density or lesser clustering of protein molecules inside the condensates of methylated FUS-eGFP. This change in the molecular packaging within these methylated FUS-eGFP condensates can be explained by the altered protein-protein interactions originating from the methylation of the positively charged arginine residues forming the cation-π interactions, the principal driver of FUS phase separation. Previous studies have shown the lesser recruitment of protein within the enhanced dynamic and liquid-like interior of methylated FUS condensates, in contrast to the unmethylated FUS-eGFP condensates^45,46^. Our anisotropy imaging readily captures this low protein density in the form of higher anisotropy values due to the diminished energy migration within these lightly packed, liquid-like condensates, as also indicated by our FRAP measurements (Fig. 4d). The picosecond time-resolved measurements (Fig. 4e) provided us with a subnanosecond component (τ_EM1_ with amplitude β_EM1_ ∼ 0.23), in addition to the intermediate and slow time constants (τ_EM2_ ∼ 1.1 ns and τ_EM3_ ∼ 9 ns) originating from the excited state energy migration within condensates. The faster energy migration rates (both τ_EM2_ and τ_EM3_) as compared to the unmethylated droplets can be attributed to the faster dynamics of the conformational fluctuations, making and breaking of contacts, and translational diffusion as a result of highly dynamic, liquid-like and reversible nature of intermolecular contacts and droplet interior within the condensates of methylated FUS.

**Fig. 4.**
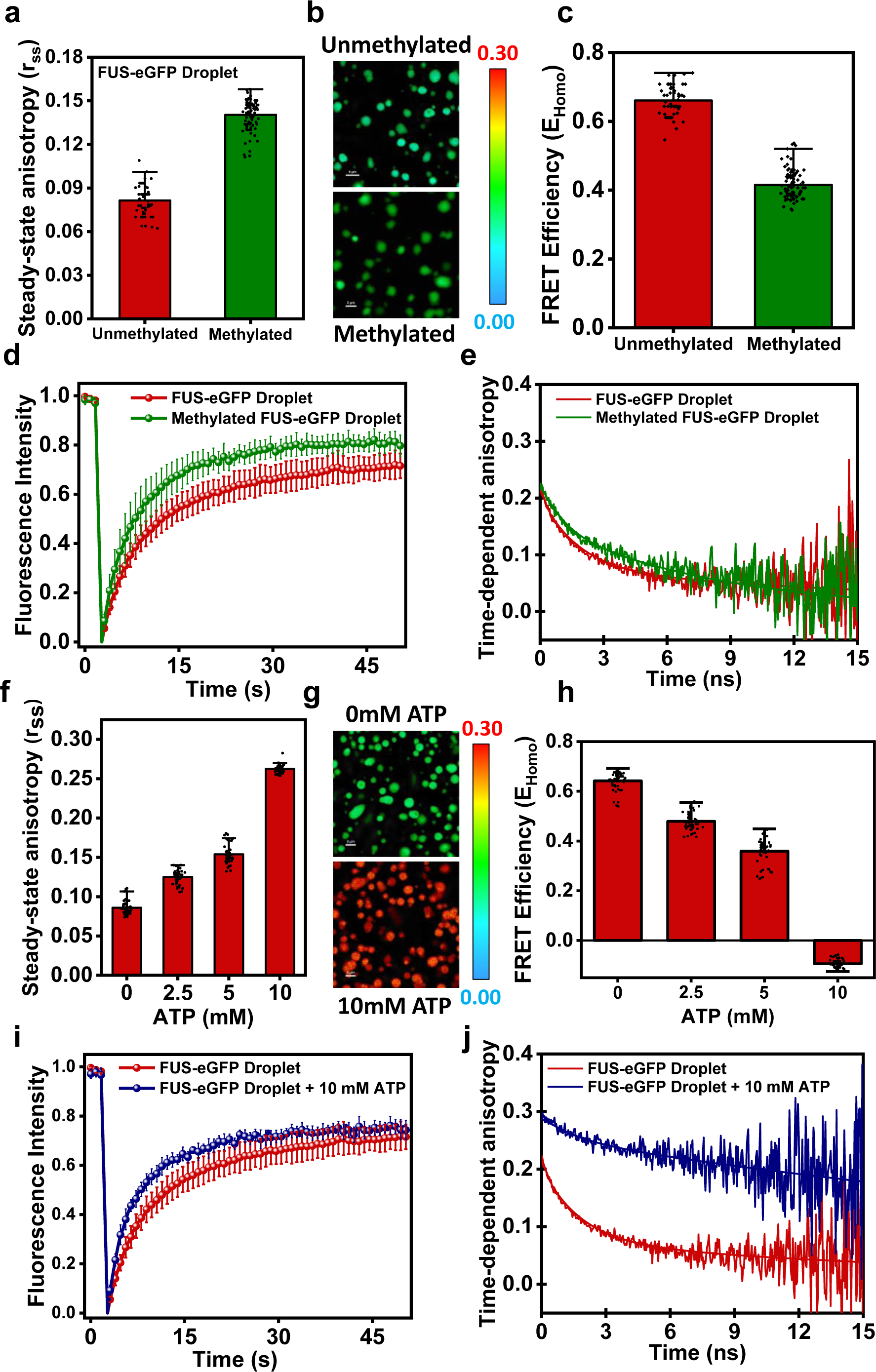
Altered molecular packing due to post-translational methylation and ATP. a. Single-droplet steady-state anisotropy plots and b. representative images obtained from unmethylated and methylated FUS-eGFP droplets. Data represent mean ± SD (n > 50 independent reactions). c. Calculated energy migration efficiencies show a reduced homoFRET efficiency within the condensates of methylated FUS-eGFP. d. Fluorescence recovery after photobleaching (FRAP) kinetics shows a faster recovery, indicating a more liquid-like dynamic interior within condensates of methylated FUS-eGFP. Data represent mean ± SD (n = 5 and 7 for unmethylated and methylated droplets, respectively) e. Representative time-resolved anisotropy decay kinetics obtained within the unmethylated and methylated FUS-eGFP droplets. f. Single-droplet steady-state anisotropy as a function of increasing ATP concentration measured within FUS-eGFP condensates. Data represent mean ± SD (n = 50 single droplets). g. Representative anisotropy images of FUS-eGFP droplets formed in the presence of 0 mM and 10 mM ATP. h. The respective FRET efficiency plot calculated from the single-droplet steady-state anisotropy values in the absence and presence of varying ATP concentrations. i. FRAP kinetics of FUS-eGFP obtained from multiple FUS-eGFP droplets in the absence and presence ATP. Data represent mean ± SD (n = 5 and 9 for 0 mM and 10 mM ATP concentration, respectively). j. Representative time-resolved anisotropy decay kinetics measured within FUS-eGFP droplets with and without ATP. Rotational correlation time and energy migration time constants are included in Table 1.

Phase separation into physiological and pathological biomolecular condensates is known to be modulated by a range of physicochemical and biochemical factors, in addition to sequence-encoded and genetic parameters. Non-peptide small molecules are being extensively studied for their ability to modulate the phase behavior and mesoscopic properties of biomolecular condensates for their potential applicability in therapeutics^47,48^. Adenosine triphosphate (ATP) is a well-known modulator of phase behavior and is shown to tune properties of biomolecular condensates in a concentration-dependent manner^49–52^. The cellular physiological concentration for ATP has been estimated to be in the range of 2-12 mM, whereas the ATP concentration of ∼ 8-10 mM is shown to cause the dissolution of protein condensates, as also in the case of FUS^49–51^. Thus, to capture the effect of ATP on the material properties of condensates through anisotropy imaging, we set up phase separation reactions of FUS-eGFP in the presence of ATP, within the physiological concentration range. Our anisotropy imaging data showed a gradual rise in anisotropy with increasing ATP concentration, and we obtained a complete recovery to the monomeric anisotropy (non-homoFRET condition) at the ATP concentration of 10 mM, above which droplet dissolution is observed (Fig. 4f, g). The dissolution effect of ATP could be monitored from the calculated homoFRET efficiency values even at the intermediate ATP concentration, where the drop in FRET efficiency suggested a gradual unpacking within condensates due to the predominance of protein-ATP interactions over the protein-protein interactions. Further, the droplets formed in the presence of 10 mM ATP exhibited no or zero-homoFRET due to the formation of extremely less dense and loosely packed condensates just prior to dissolution (Fig. 4h). This dynamic, liquid-like droplet interior just prior to dissolution was also evident from the fast and complete FRAP recovery obtained with 10 mM ATP FUS-eGFP droplets (Fig. 4i). Our time-domain fluorescence anisotropy measurements obtained within these droplets yielded biexponential decay kinetics with a minimal contribution of the intermediate energy migration time constant (τ_EM2_ ∼ 1.6 ns) and a major contribution from the rotational correlation time component (ϕ ∼ 50 ns) (Fig. 4j). This relatively faster rotational diffusion also suggests the enhanced diffusivity and liquid-like interior of the FUS-eGFP droplets formed in 10 mM ATP, as compared to the FUS droplets formed in the absence of ATP (ϕ ∼ 60-70 ns) (Fig. 2j). After evaluating the sensitivity of homoFRET imaging for condensates formed under a wide range of solution conditions, we next aimed to utilize homoFRET to study cellular phase-separated assemblies.

### Mapping intermolecular organization within phase-separated assemblies of FUS *in situ*

Fluorescently tagging proteins of interest with fluorescent proteins such as GFP, CFP, YFP, mCherry, etc., is a standard practice employed for cellular studies investigating the localization, conformational changes, protein-protein association, and so on. The majority of these fluorescent proteins exhibit a significant spectral overlap between their own absorption and emission spectra, making them a suitable candidate for such homoFRET studies. Thus, we next asked whether anisotropy imaging could be employed to detect the formation of these dynamic liquid-like assemblies within the crowded, complex cellular milieu. To explore this idea, we began with the transient expression of eGFP-tagged wild-type FUS in human lung epithelial cell line, A549. The amino acid sequence of FUS harbors a nuclear localization signal (NLS) at the C-term end (Fig. 2a) and, hence, primarily resides in the nucleus, where it is involved in functions such as genome organization, DNA damage repair, transcription, RNA splicing, translational regulation, and so on^36,37,44^. Upon overexpression in the cells, FUS readily localized to the nucleus and was more or less uniformly distributed throughout the nucleoplasm, as seen in our confocal microscopy imaging (Fig. 5a). However, a significant proportion of the cells also showed the formation of small nuclear puncta or foci, presumably as a consequence of the overexpression stress within cells. To probe the distinct molecular packaging and distribution of FUS within the dispersed nucleoplasm and condensed foci, we began with anisotropy imaging of these cells overexpressing wild-type eGFP-FUS (Fig. 5b, c).

**Fig. 5.**
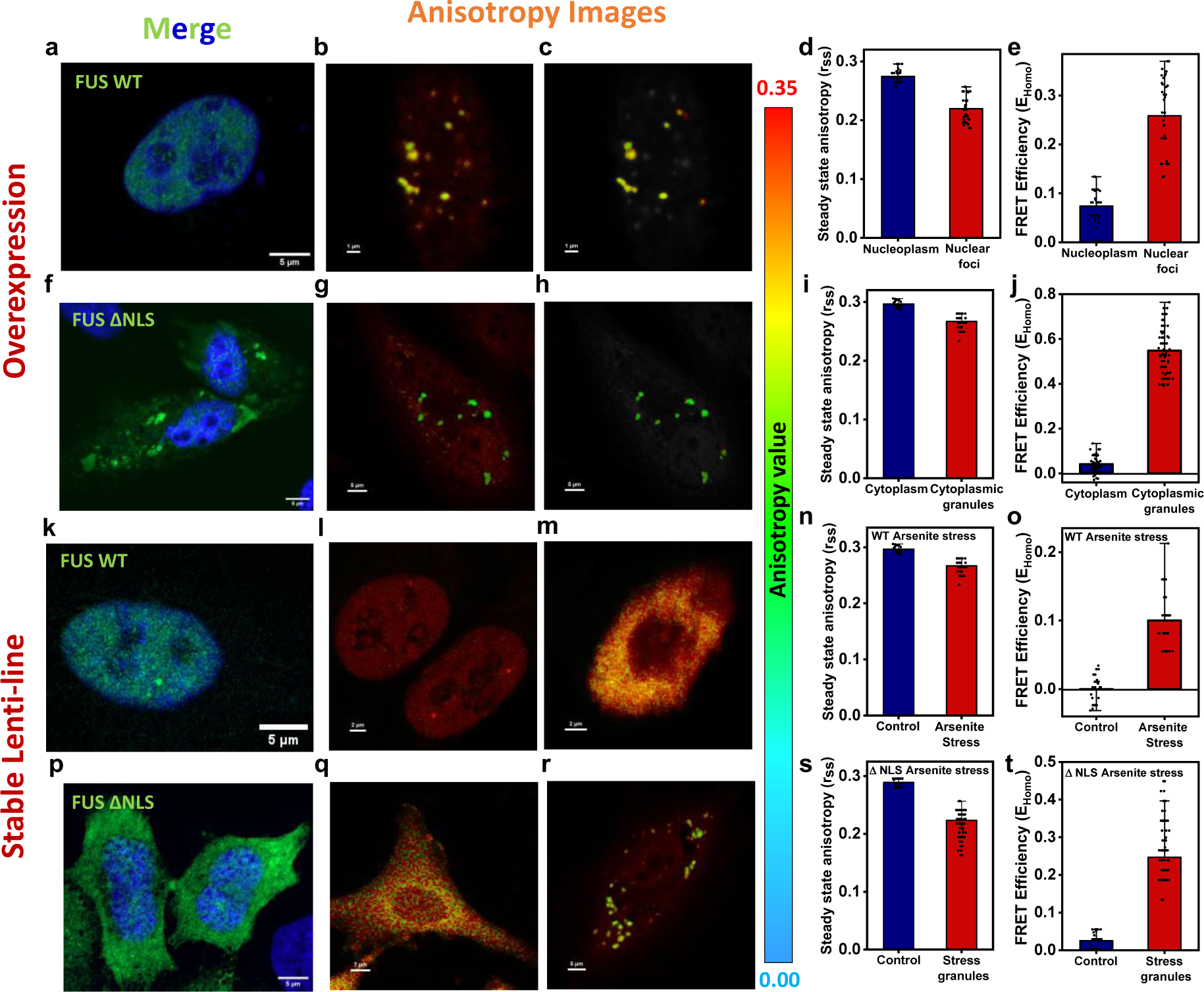
Molecular packing within phase-separated assemblies of FUS *in situ*. a. Representative confocal airyscan image of A549 cells overexpressing wild-type eGFP-FUS (green) show diffused localization of eGFP-FUS inside the nucleus. b. Anisotropy images of eGFP-FUS overexpression highlighting the nuclear foci (c). d. Steady-state anisotropy plots obtained from the diffused nucleoplasm and condensed nuclear foci with corresponding FRET efficiencies (e). Data represent mean ± SD (n > 30 measurements). f. Representative confocal airyscan image of A549 cells transiently expressing eGFP-FUS ΔNLS mutant showing cytoplasmic localization along with recruitment into cytoplasmic granules. g. Representative anisotropy image of cells transiently expressing mutant eGFP-FUS indicating a loss in anisotropy within the cytoplasmic granules (h), (i). j. HomoFRET efficiency plot showing energy migration efficiency within the diffused cytoplasm and cytoplasmic granules. Data represent mean ± SD (n ≥ 30 measurements). k. Representative airyscan confocal image of A549 cells with stable expression of wild-type eGFP-FUS with nuclear localization.

Representative anisotropy images of cell nuclei in the absence (l) and presence of (m) arsenite stress. Cells stably expressing eGFP-FUS WT and eGFP-FUS ΔNLS were treated with 300 µM sodium arsenite for 1 h to induce stress granule formation. n. Steady-state anisotropy plot and the corresponding FRET efficiency plot (o), obtained from the nucleus of cells expressing wild-type eGFP-FUS in the non-stressed and stressed conditions. Data represent mean ± SD (n ≥ 30 measurements). p. Airycan confocal image of A549 cells with stable eGFP-FUS ΔNLS expression showing mislocalization of mutant eGFP-FUS into the cytoplasm, without the formation of cytoplasmic granules. Representative anisotropy images of cells expressing the eGFP-FUS ΔNLS mutant showing q. diffused in non-stressed conditions and r. cytoplasmic stress granules upon exposure to arsenite stress. Corresponding steady-state anisotropy plot (s) and FRET efficiency plot (t) comparing the non-stressed and arsenite stress conditions. Data represent mean ± SD (n > 40 measurements).

In line with our hypothesis, the steady-state anisotropy value showed a significant change in the nucleoplasm and foci comprising eGFP-FUS (Fig. 5d). The loss in anisotropy within the nuclear foci suggested clear compaction and denser packaging as indicated by the more than 3-fold increase in the apparent energy migration efficiencies, as compared to surrounding dispersed phase (Fig. 5e). As mentioned previously, FUS is associated with the pathology of ALS, where a significant fraction of familial ALS cases is attributed to point mutations within FUS NLS, resulting in the accumulation of insoluble pathological aggregates due to cytoplasmic mislocalization^38,45,53–55^. Thus, with the objective of recapitulating the ALS-associated cytoplasmic mislocalization, we next overexpressed a ΔNLS mutant of FUS (FUS ΔNLS) in these cells to visualize the cytoplasmically localized eGFP-FUS. Our fluorescence microscopy imaging revealed the mislocalization of FUS in the cytoplasm, in addition to the formation of cytoplasmic granules, as a response to the overexpression stress (Fig. 5f). Our steady-state anisotropy imaging captured a drastic change in the molecular packaging within the concentrated granules and the cytoplasmically dispersed FUS (Fig. 5g-i). The calculated FRET efficiencies exhibited nearly 13-fold increase in the excited state energy migration, capturing the reduced intermolecular distances and enhanced protein-protein associations in the cytoplasmic puncta (Fig. 5j). Next to probe the molecular packing within the phase-separated nuclear and cytoplasmic assemblies of FUS, we began with anisotropy imaging of cells with stable expression of wild-type and mutant FUS.

One of the well-studied aspects of FUS physiology is its assembly and recruitment of stress granules in response to stress conditions. Upon exposure to various kinds of stress, cells attempt to protect themselves from stress-related damage and death by halting the ongoing energy-consuming processes, including transcription and translation. Stress granules provide a sink for the sequestration of translationally stalled mRNAs, ribosomes, translation initiation, and elongation factors, along with several other RNA-binding proteins in the form of membraneless, dynamic, liquid-like RNP granules to facilitate cell survival. We next sought to capture this altered localization, packaging, and protein-protein interaction of eGFP-FUS within the liquid-like phase-separated stress granules assembled in response to cellular stress. With this objective, we began with the stable expression of wild-type FUS and FUS ΔNLS. To invoke a stress damage response, we subjected the cells to arsenite exposure, which has previously been shown to induce FUS-containing stress granule formation within cells^55–58^. Confocal fluorescence imaging revealed completely diffused eGFP-FUS in the nucleus in the absence of arsenite stress and correspondingly high anisotropy due to the non-compact packaging of eGFP-FUS within the healthy nucleus (Fig. 5k). Upon incubation with sodium arsenite, cells expressing wild-type protein exhibited nuclei with diffused FUS, however, with a relatively denser packaging, presumably due to enhanced nuclear oligomerization owing to arsenite-induced stress response (Fig. 5l, m). Steady-state anisotropy measurements provided a slightly reduced anisotropy value, suggesting a relatively higher density or protein-protein association within the nuclei of stressed cells, leading to a larger energy migration efficiency (Fig. 5n, o). Lastly, we sought to probe the organization and packaging inside stress granules formed within cells with stable expression of the FUS ΔNLS mutant. The mutant FUS exhibited extensive mislocalization into the cytoplasm, which was further recruited into phase-separated stress granules in response to arsenite stress (Fig. 5p-r). Comparative anisotropy measurements revealed an approximately 6-fold increase in the extent of energy migration reporting on the closely packed architecture and intermolecular interactions within the stress granules formed upon arsenite stress (Fig. 5s, t). Taken together, our data underscore the efficacy of homoFRET imaging as a direct readout for condensate formation and further investigations into the interior organization of supramolecular protein assemblies formed *in situ* via phase separation.

## Discussion

Unique supramolecular assemblies of proteins and nucleic acids formed via the process of biomolecular condensation carry great physiological and pathological significance across organisms. The prime distinguishing factor between the majority of these functional, phase-separated, reversible condensates and irreversible pathological deposits or aggregates that are associated with aging is their internal material properties. These characteristics are determined by the internal organization and intermolecular interactions within these assemblies. Our work demonstrates the utilization of homoFRET, as a potent tool for discerning the material characteristics of condensates formed *in vitro* and *in situ* while assessing the extent of molecular packaging and intermolecular association quantitatively. By performing anisotropy imaging, we obtained steady-state anisotropy values as a measure of excitation energy migration via homoFRET. Energy migration from the excited state of one fluorophore to a chemically identical, proximally placed fluorophore leads to a depolarization of the emitted fluorescence, which is measured by the loss in the anisotropy. The extent of energy transfer via homoFRET depends on the clustering density or the higher-order packing of the fluorophore molecules, which can be readily determined from the magnitude of emission depolarization. In this work, we demonstrate the utility of anisotropy imaging to elucidate the dynamic internal architecture and obtain a quantitative measure of molecular proximities (E_homoFRET_) within biomolecular condensates of a neuronal protein FUS, formed in the absence and presence of multiple phase separation modulators *in vitro* and *in situ*.

Fused in Sarcoma (FUS) is an RNA-binding protein with a prion-like low-complexity domain associated with various functional and pathological aspects and known to undergo phase separation within cells^31–33,45,46^. Anisotropy imaging of FUS tagged with eGFP resulted in a sharp dip in the steady-state anisotropy upon phase separation, corresponding to the multiple-fold increase in energy migration efficiency due to the extensive molecular crowding and compact packaging in the demixed or condensed phase. This anisotropy loss could be completely recovered upon photobleaching of the droplets, verifying homoFRET as the origin of emission depolarization. Further, our picosecond time-resolved measurements yielded mono and biexponential decay kinetics, depending on the contribution of homoFRET towards fluorescence depolarization. The depolarization decay in homoFRET conditions could be further differentiated into multiple energy migration components and rotational dynamics of FUS-eGFP with their corresponding time constants and amplitudes. These energy migration rates provide a readout for the diverse molecular level events and their distinct timescales contributing to the multiple energy migration modes facilitating homoFRET. Steady-state anisotropy values exhibited remarkable sensitivity to the variation in local densities of FUS-eGFP, as evident in our ratiometric droplet measurements. Our anisotropy imaging also captured the effect of RNA on the droplet architecture and higher-order packaging of FUS condensates, doped with FUS-eGFP, as a homoFRET reporter. At lower concentrations, RNA enhances the protein-protein association, as shown previously, leading to a relatively dense droplet interior with increasing homoFRET efficiency. At higher concentrations, just prior to dissolution, the droplets exhibit a heterogeneous organization reminiscent of the core-shell morphology previously shown for other phase-separating systems. The core showed comparatively higher molecular proximities (high E_homoFRET_) corresponding to the visibly low steady-state anisotropy values with respect to the lightly-packed periphery. This was followed by the sudden rise in anisotropy values to that of the dispersed monomeric protein upon dissolution of the condensates. We also captured the effect of post-translational methylation and small-molecule phase separation modulator, namely ATP, on the internal packaging and droplet properties of FUS-eGFP. Methylation resulted in sparsely packed droplets with a more liquid-like interior, suggesting diminished clustering and intermolecular interactions in comparison to the unmethylated FUS-eGFP condensates. Similarly, with increasing ATP concentration, anisotropy values showed a gradual recovery to that of the non-homoFRET conditions, in agreement with the previously established dissolution effect of ATP^49,50^. This unpacking of FUS-eGFP condensates upon methylation and in the presence of ATP can be ascribed to the altered propensity of arginine to associate with the N-terminal tyrosine residues via cation-π interactions. Lastly, we utilized our methodology to investigate the condensate formation and higher-order packing within phase-separated assemblies of FUS-eGFP in mammalian cell lines. Our results highlight the altered densities and molecular packing within these phase-separated condensates formed under divergent cellular conditions and localization. Taken together, our data showcases the sensitive technique of anisotropy imaging as a versatile tool to probe the fundamental property of molecular packaging and the internal architecture of biomolecular condensates formed via phase separation. Cellular studies that regularly employ tagging of molecules with fluorescent proteins can be readily extended to gain deeper insights into the organization and supramolecular packing within the dynamic membraneless cellular compartments. A wide range of fluorophores, including the extensively used fluorescent proteins (RFP, YFP, CFP, mCherry, etc.), can be utilized for homoFRET imaging within condensates both *in vitro* and *in situ*. In addition to phase separation, homoFRET via anisotropy imaging can also report on the higher-order species or nanoclusters formed as the precursors of these supramolecular assemblies^59,60^. Previous studies have highlighted the use of fluorescent protein lifetimes as a measure of droplet densities and crowding^61^. In addition to the droplet interior, steady-state anisotropy can distinctly report on the alterations in polypeptide chain clustering and condensate architecture resulting from the intermolecular associations characteristic of the given condensates. The requirement of a single fluorescent probe for homoFRET broadens the scope substantially and, thus, the availability of fluorescent reporters for simultaneously tagging multiple proteins of interest. HomoFRET

studies with two or more homoFRET reporters in conjunction with intra or intermolecular heteroFRET studies can further shed light on the critical molecular events associated with multicomponent heterotypic phase separation^62–64^. Previous studies have utilized homoFRET to identify and characterize membrane-induced clustering and oligomerization *in vitro* as well as in membrane-anchored proteins *in situ*^65^. HomoFRET imaging can be employed to investigate membrane-associated phase separation of membrane-bound proteins^66,67^. Recent studies have shown the tuning of FUS condensates and their material properties by the protein quality control (PQC) machinery, including small heat shock proteins and chaperones^68,69^. HomoFRET imaging can be successfully employed to uncover the molecular details and reorganization associated with this chaperoning effect of the PQC machinery on the preformed condensates of FUS and other pathological proteins. In contrast to the original hypothesis, several reports now suggest the presence of a heterogeneous organization or a distinct small-world architecture within biomolecular condensates^41–43,59,70^. Anisotropy imaging can illuminate this spatial heterogeneity, presumably arising due to the non-uniform, selective distribution and interactions of constituent proteins and nucleic acids within the phase-separated droplets. These localized density variations will be efficiently recorded in the homoFRET efficiency fluctuations in the form of distinct steady-state anisotropy values throughout the condensates. Thus, we believe this methodology can provide a unique approach to discern the condensate packaging and differential intermolecular interactions within a wide range of biological condensates and facilitate a comprehensive understanding of *in vitro* and cellular phase-separated assemblies.

## Methods

### Recombinant protein expression and purification

The plasmids pMal-*Tev*-FUS-*Tev*-His_6_ and pMal-*Tev*-FUS-EGFP-*Tev*-His_6_ were transformed into *E. coli* BL21(DE3) RIPL bacterial strain, for overexpressing MBP-*Tev-*FUS-*Tev-* His_6_ (referred to as FUS hereafter) and MBP-*Tev-*FUS-EGFP-*Tev-*His_6_ (referred as FUS-eGFP hereafter) respectively. Overexpressed recombinant FUS and FUS-eGFP were purified by using tandem Ni-NTA and amylose resin affinity chromatography. For FUS and FUS-eGFP overexpression, bacterial cultures were grown in LB media at 37 °C, 220 rpm till an O.D._600_ of 0.6–0.8, and protein expression was induced by adding 0.1 mM isopropyl-β-thiogalactopyranoside (IPTG) at 12 °C, 220 rpm for 22 h. Bacterial cells were harvested by centrifuging at 4 °C, 3220 × *g* for 40 min. Cell pellets were stored at −80 °C for further use.

For purification, pellets were resuspended in lysis buffer (50 mM sodium phosphate, 300 mM NaCl, 40 mM imidazole, 10 μM ZnCl_2_, 4 mM BME, and 10% v/v glycerol, pH 8.0), and bacterial cells were lysed by probe sonication at 5% amplitude, 15 s ON and 10 s OFF for 25 minutes. This bacterial whole-cell lysate was centrifuged at 4 °C, 15,557 × *g* for 1 h, to obtain the supernatant, which was then incubated with lysis buffer equilibrated Ni-NTA agarose beads for 1.5 h at 4 °C. The beads were washed with wash buffer and protein was eluted with 250 mM imidazole, followed by binding to the amylose resin. Protein was eluted from amylose resin with 20 mM maltose elution buffer (50 mM sodium phosphate, 800 mM NaCl, 40 mM imidazole, 10 μM ZnCl_2_, 20 mM maltose, and 1 mM 1,4-dithiothreitol, pH 8.0). The concentration of FUS and FUS-eGFP was estimated by measuring absorbance at 280 nm (using ɛ_280_ calculated from the Scripps Protein Calculator v3.4). Purified protein samples were run on SDS-PAGE gel to check protein purity. Purified proteins were temporarily stored at 4 °C and freshly concentrated for *in vitro* phase separation assays.

The pET28b-PRMT1 plasmid was transformed into *E. coli* BL21(DE3) std bacterial strain, for overexpressing His_6_-PRMT1. For overexpression, bacterial cultures were grown in LB media till O.D._600_ of 1, and protein expression was induced by adding 1 mM isopropyl-β-thiogalactopyranoside (IPTG) at 20°C, 220 rpm for 16 h. Bacterial cells were harvested by centrifuging at 4 °C, 3220 × *g* for 40 min, and pellets were stored at −80 °C for further use. For protein purification, pellets were resuspended in lysis buffer (50mM Tris-HCl pH 7.5, 150 mM NaCl, 20 mM imidazole, 4 mM BME, 20% glycerol), and bacterial cells were lysed by probe sonication at 5% amplitude, 15 s ON and 10 s OFF for 25 minutes. The cell lysate was centrifuged at 4 °C, 15,557 × *g* for 1 h, for separation of the insoluble cellular debris. The supernatant was passed through pre-equilibrated Ni-NTA agarose beads. The beads were washed with a gradient of imidazole, and the protein was eluted in elution buffer (50 mM Tris-HCl pH 7.5, 1 M NaCl, 500 mM imidazole, 1 mM DTT). Protein was further purified and buffer exchanged by size exclusion chromatography using HiLoad 16/600 Superdex-G-200 (GE Healthcare) gel filtration column to storage buffer (50mM Tris-HCl pH 7.5, 150 mM NaCl, 20 mM imidazole, 4 mM BME, 20% glycerol) and stored at −80 °C for future use.

### Phase separation assays

Phase separation of FUS and FUS-eGFP was induced by the addition of TEV protease (TEV: protein molar ratio of 1:10) in the phase separation buffer (20 mM HEPES, 1 mM DTT, pH 7.4) and incubated at room temperature for 10 min. For all phase separation experiments, total protein concentration was fixed at 10 μM, and for ratiometric measurements, the fraction of FUS-eGFP was varied from 1% to 100% (FUS-eGFP only droplet). For RNA-dependent anisotropy measurements, phase separation of 10 μM FUS (2 μM FUS-eGFP + 8 μM FUS) was set up in the presence of varying RNA concentrations (0 ng/μL, 25 ng/μL, 50 ng/μL, 75 ng/ μL, and 100 ng/μL polyU). To monitor the effect of methylation, phase separation of methylated FUS-eGFP was set up at a protein concentration of 10 μM for anisotropy measurements. For phase separation in the presence of ATP, FUS-eGFP droplet formation was induced in a buffer supplemented with MgCl_2_ in the presence of 0 mM, 2.5 mM, 5 mM, and 10 mM ATP.

### Steady-state anisotropy imaging

Anisotropy imaging was performed using our confocal-based MicroTime 200 time-resolved microscope to obtain single-droplet steady-state anisotropy values. A pulsed laser (485 nm, 20 MHz) and a Super Apochromat water immersion 60x objective (Olympus, NA 1.2) were used for performing anisotropy measurements in the dispersed and droplet phases. Droplet formation was induced under varying solution conditions (RNA, ATP, methylation) and incubated at room temperature for 10 min. Droplet reactions were spotted on a glass coverslip and allowed to settle on the surface. A549 cells with transient and stable expression of eGFP-FUS and eGFP-FUS ΔNLS were separately grown on glass coverslips and fixed with 4% w/v paraformaldehyde (PFA) in PBS buffer at room temperature (RT) for 10 min, prior to anisotropy imaging. Upon excitation with the laser, the emitted fluorescence was split into two separate channels using a polarizing beam-splitter and directed to the single-photon counting avalanche diodes (SPADs). Anisotropy images were generated from the parallel and perpendicular counts after incorporating the correction factors using SymphoTime64 software v2.7. The steady-state anisotropy was calculated using eq^n^ (1) as follows;

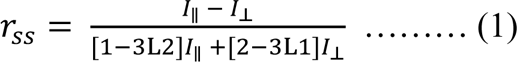

where *I*_∥_ and *I*_⊥_ are the parallel and perpendicular fluorescence intensities after background correction, and L1 (0.0308) and L2 (0.0368) account for the objective correction factors.

Based on the steady-state anisotropy values, energy migration efficiencies (E_homoFRET_) within the condensates were calculated using eq^n^ (2) as follows^26^;

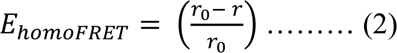

where r_0_ and r are the calculated steady-state anisotropy values in the absence and presence of homoFRET. For FRET efficiency calculations, we used the monomeric steady-state anisotropy (non-homoFRET condition) value as the r_0_.

### Time-resolved anisotropy measurements

For the picosecond time-resolved fluorescence anisotropy measurements, freshly phase-separated FUS-eGFP droplets were settled, and fluorescence intensity was obtained in a droplet-by-droplet manner by focusing the pulsed laser inside individual droplets. The parallel and perpendicular intensity decay profiles were obtained and extracted using SymphoTime64 software v2.7 for further decay analysis to perform global fitting using eqn (3) and (4),

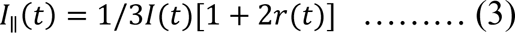

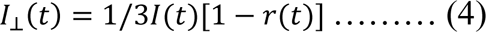

where *I*_||_(t), *I*_⊥_(t), and *I*(*t*) can be defined as the time-dependent fluorescence intensities collected at the parallel, perpendicular, and magic angle (54.7°) geometry. The G-factor correction was incorporated for the perpendicular decay component, estimated by collecting the fluorescence intensity of free dye in the parallel and perpendicular channels using the anisotropy setup. The picosecond time-resolved fluorescence anisotropy decays were fitted using a suitable decay model (mono/bi/triexponential) based on the goodness of fit estimated from the autocorrelation function, randomness of residuals, and reduced χ2 values^71^. The anisotropy decay kinetics were used to obtain energy migration time constants (τ_EM_) and rotational correlation time (ϕ) as follows;

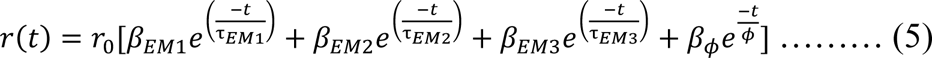

where r_0_ denotes the (time-zero) fundamental anisotropy of eGFP, and β_EM1_, β_EM2_, β_EM3_, and β_ϕ_ are the amplitudes associated with the rotational correlation time (ϕ), fast (τ_EM1_), intermediate (τ_EM2_), and slow (τ_EM3_) energy migration time constants, respectively.

The amplitude for ultrafast energy migration component (β_EM1_) was estimated based on the unresolved (< 100 ps) loss in the time-zero anisotropy (r_0_) in homoFRET conditions, as follows;

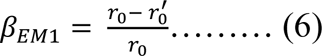

where, r_0_ is the fundamental anisotropy and r_0_’ is the time zero-anisotropy in the presence of energy migration via homoFRET.

### Time-dependent photobleaching anisotropy imaging

Phase separation of 10 µM FUS-eGFP was set up and incubated at room temperature for 10 min. Droplet solution (∼ 50 µL) was spotted on the glass coverslip, and anisotropy imaging was performed at maximum laser power at fixed time intervals for up to 10 min. Steady-state anisotropy values were obtained for three independent reactions. Data represent the mean and standard deviation (n = 3).

### *In vitro* methylation

Prior to setting up the methylation reaction, PRMT1 was buffer exchanged to the *in vitro* methylation buffer (IVM) (50 mM Na_2_HPO_4_, 150 mM NaCl, 5% glycerol, 1 mM EDTA, 1 mM DTT, pH 8.0) using a PD-10 column. Pure PRMT1 was concentrated using 10 KDa MWCO Amicon filters and used for the methylation of FUS-eGFP. Freshly purified FUS-eGFP was concentrated and washed with the IVM buffer prior to methylation. The methylation reaction was set up at a molar ratio of 1:2 (FUS-eGFP: PRMT1) in the presence of 1mM S-adenosyl-L-methionine (SAM) as the methyl donor in the IVM buffer and incubated at room temperature overnight. Methylated FUS-eGFP was concentrated freshly for droplet anisotropy measurements.

### Fluorescence Recovery after Photobleaching (FRAP) measurements

Phase separation of 10 µM FUS-eGFP was induced by the addition of TEV in the phase separation buffer (20 mM HEPES, 1 mM DTT, pH 7.4). Reactions were incubated at room temperature for 10 min, and a 10 µL sample was placed on a glass coverslip. FRAP experiments were performed on the ZEISS LSM 980 instrument using a 63× oil-immersion objective (NA 1.4). Droplets were allowed to settle, following which a region of diameter 1 µm was selected and bleached with the help of the green laser (488 nm laser diode) and the fluorescence was monitored with time using a monochrome cooled high-resolution AxioCamMRm Rev. 3 FireWire(D) camera. The fluorescence recovery was recorded with the Zen Blue 3.2 (ZEISS) software, corrected for the background fluorescence, and plotted using the Origin software. FRAP measurements were obtained within the droplets of FUS-eGFP formed in the absence and presence of ATP and polyU RNA.

### Cell culture and confocal imaging

A549 (obtained from ATCC) were grown in DMEM High Glucose (Gibco) supplemented with 10% FBS Heat-Inactivated (Gibco), GlutaMAX (Gibco), sodium pyruvate (Gibco), MEM NEAA (Non-Essential Amino Acids – Gibco), penicillin, streptomycin, and glutamine (Gibco), in a humidified incubator at 37°C, with 5% CO_2_. For overexpression, A549 cells were transiently transfected with FUS-pEGFP C1 or FUS ΔNLS-pEGFP C1 using Lipofectamine LTX with PLUS Reagent (Thermo Fisher Scientific). A549 cells stably expressing eGFP-FUS (1 to 526), and eGFP-FUS ΔNLS (1 to 514) were generated by lentiviral transduction. Transduced A549 cells stably expressing desirable levels of eGFP-FUS, eGFP-FUS ΔNLS were sorted using BD FACS Aria™ Cell Sorter and expression of eGFP-FUS and eGFP-FUS ΔNLS was verified by immunofluorescence. For fluorescence confocal imaging, cells were grown on glass coverslips and fixed with 4% w/v paraformaldehyde (PFA) in PBS buffer at room temperature (RT) for 10 min. After fixation, nucleus was stained with, followed by mounting on glass slides using Fluoromount-G™ Mounting Medium.

Fluorescence confocal microscopic imaging of fixed A549 cells expressing eGFP-FUS or eGFP-FUS ΔNLS, with or without arsenite stress, counterstained with DAPI, was performed using ZEISS LSM 980 using a 63× (NA 1.4) oil-immersion objective, with 408 nm (DAPI) and 488 nm (eGFP) laser diode. The images were acquired at 1024 × 1024 pixels and 16-bit depth resolution, with 2× averaging, by a monochrome-cooled high-resolution AxioCamMRm Rev. 3 FireWire(D) camera. Image processing and analyses were performed on in-built Zen Blue 3.2 software and ImageJ (NIH, Bethesda, USA).

### Sodium arsenite treatment

A549 cells stably expressing eGFP-FUS or eGFP-FUS ΔNLS were treated with 300 µM sodium arsenite for 1 hour, to induce stress granule (SG) formation. After treatment, cells were fixed, stained with DAPI and mounted on glass slides using Fluoromount-G™ Mounting Medium for confocal microscopic imaging.

## Acknowledgments

We thank IISER Mohali, Science and Engineering Research Board (SUPRA SPR/2020/000333 to S.M.), Department of Science and Technology, Govt. of India (FIST grant # SR/FST/LS-II/2017/97 to the Department of Biological Sciences, IISER Mohali), and Ministry of Education, Govt. of India (Centre of Excellence grant to S.M. and the Prime Minister’s Research Fellowship to A.W. and D.C.) for financial support. We thank the Indo-French Centre for the Promotion of Advanced Research (IFCPAR/CEFIPRA) for fellowship to L.A. We thank Prof. Dorothee Dormann (Johannes Gutenberg University of Mainz, Germany) for her kind gift of FUS full-length, FUS-eGFP, and PRMT1 plasmids. Dr. Tatyana Shelkovnikova (Sheffield Institute for Translational Neuroscience (SITraN), The University of Sheffield, UK) for her kind gift of FUS-pEGFP C1 and FUS ΔNLS-pEGFP C1 plasmids. We thank Professor N. Periasamy (Retd. TIFR Mumbai) for the fluorescence decay analysis program.

## Authors contributions

A.J. and S.M. conceived the project. A.J., A.W., and S.M. further developed the concept and the experimental design. A.J., A.W., S.S., L.A., and D.C. performed the experiments and analyses. G.K., P.J., and I.B. provided cells with transient and stable expression of wild-type and mutant FUS. A.J. and A.W. prepared the figures and wrote the first draft. S.M. supervised the work, wrote/edited the manuscript, obtained funding, and provided the overall direction. All authors discussed the results and commented on the manuscript.

